# Computational study of heme *b*_595_ to heme *d* electron transfer in E. coli cytochrome *bd*-I oxidase

**DOI:** 10.1101/2025.09.03.673948

**Authors:** Raaif Siddeeque, Baptiste Etcheverry, Côme Cattin, Jean Deviers, Frédéric Melin, Petra Hellwig, Fabien Cailliez, Aurélien de la Lande

## Abstract

Cytochrome *bd* is a distinctive family of terminal oxidases present in the respiratory chains of many prokaryotes. Despite its biological importance, the redox chemistry of these proteins remains poorly understood, largely due to the presence of two *b*-type hemes and one *d*-type heme. Here, we report the first computational study of inter-heme electron transfer in the cytochrome *bd* family. We performed 10 μs of molecular dynamics simulations of *E. coli* cytochrome *bd*-I embedded in realistic membranes, combined with quantum chemical calculations to estimate the thermodynamic parameters of electron transfer from heme *b*_595_ to heme *d* within the framework of Marcus theory. We further identify the respective contributions of the hemes, protein scaffold, lipid bilayer, water, and counterions to the driving force and reorganization energy. The inter-heme electronic coupling was calculated using the Projected Orbital Diabatization (POD) method in a hybrid Quantum Mechanics/Molecular Mechanics scheme and rationalized through electron transfer pathway analysis. This study provides fundamental insights into how electron transfer steps are orchestrated in the catalytic cycle of *E. coli* cytochrome *bd*-I.

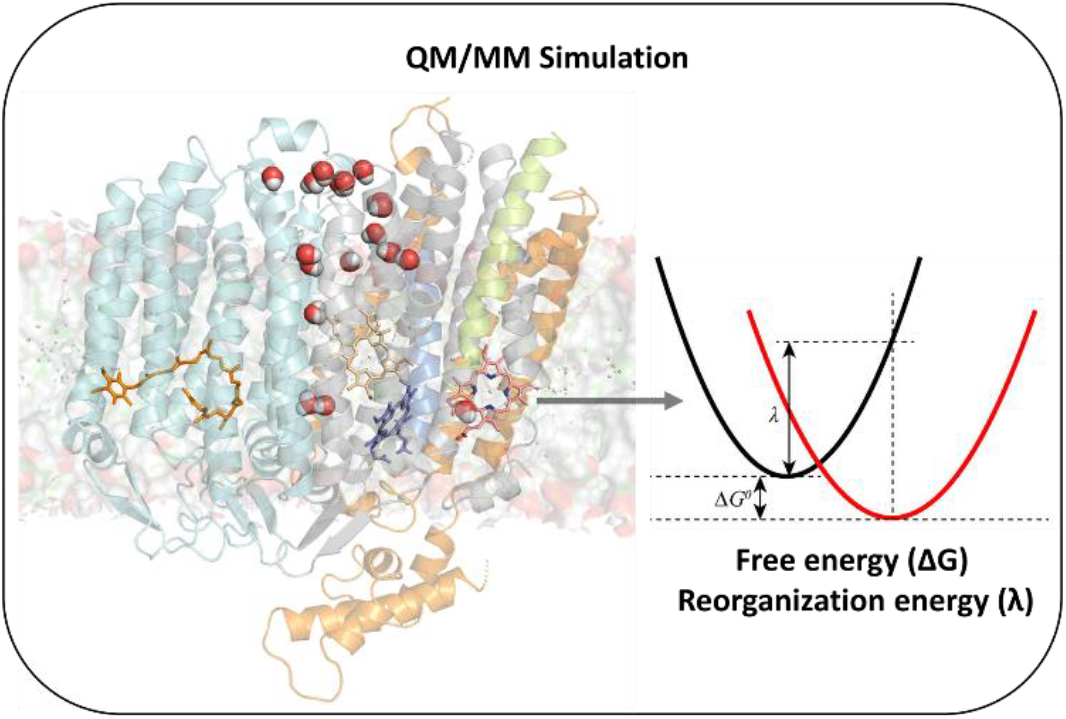

## 1. Introduction

Oxygen reduction is a crucial step in the aerobic respiration. Three enzyme superfamilies are able to catalyze oxygen reduction to water, including *bd* oxidases. These highly diverse terminal oxidases can be found in both archaea and bacteria [1–3]. Well-characterized *bd* oxidases are found in *E. coli*, where at least one quinol, two *b*-type hemes, and one *d*-type heme participate in electron transfer steps during the catalytic process. Molecular oxygen is reduced at the *d* heme. As described by the recent structural studies on *bd* oxidases from different organisms [3] and by evolutionary studies [2], a high degree of diversity was recently identified. At present the X-Ray structure of the cytochrome *bd* oxidase from *Geobacillus thermodenitrificans* [4] and the Cryo EM structures of those from *E. coli* [5–7], *Corynebacterium glutamicum* [8], *Mycobacterium smegmatis* [9] and *Mycobacterium tuberculosis* [10] have been described. These structures reveal that the position of the hemes is dependent on the organism. Electrochemical studies also illustrate that the heme potential is strongly variable among *bd* oxidases, pointing to a major adaptation of these enzymes to the environment the organism (bacteria or archaea) is exposed to. More specifically, in cyt *bd*-I from *E. coli*, redox potentials of +258 mV for heme *d*, +176 mV for heme *b*_558_ and +168 mV for heme *b*_595_ were reported [11]. In *bd-*II upshifted potential values were found for all hemes [7]. However, in *Geo. th* values of +155 mV for heme *b*_558_, +50 mV for heme *b*_595_ and +15 mV for heme *d*, were found [6]. In order to get insight into the role of the structure for the electron transfer properties in *bd* oxidases, molecular simulation is a powerful tool. In this paper, we use a combination of molecular dynamics simulations and quantum chemistry calculations to study electron transfer in cyt *bd*-I from *E. coli*.

The rather uncommon hemes’ coordination sphere (Figure 1) certainly contribute to fine tune O_2_ reduction catalysis, although the details of the reaction mechanism remain unclear. On one hand, heme *b*_558_ is coordinated to Met393 and His186. It is assumed to be the first acceptor for the electrons coming from quinol oxidation. On the other hand, heme *b*_595_ is likely to be coordinated by the side chain of Glu445 in *E. coli* [6]. The protonation state of the latter residue is unknown, and could actually vary during the catalytic cycle to compensate charge variations within the active site caused by proton or electron transfers [6]. It was shown that Glu445 mutation by a non-ionizable residue (Ala) made heme *b*_595_ more difficult to reduce [12]. At that time however, it was not known that Glu445 was a heme *b*_595_’s ligand. The redox potentials of the Glu445Ala variant were similar to those of the wild type [13], whereas they were significantly upshifted in the Glu445Asp variant, as well as in a mutant of the Arg448 which is susceptible to form a salt bridge with Glu 445 [14]. Finally, these proteins encapsulate a heme *d* which contains only one propionate function and a spiro-lactone chemical group. We note that both heme *b*_595_ and heme *d* leaves the possibility for a sixth coordination, by external ligands (*e.g*. O_2_, H_2_O).

**Figure 1:**
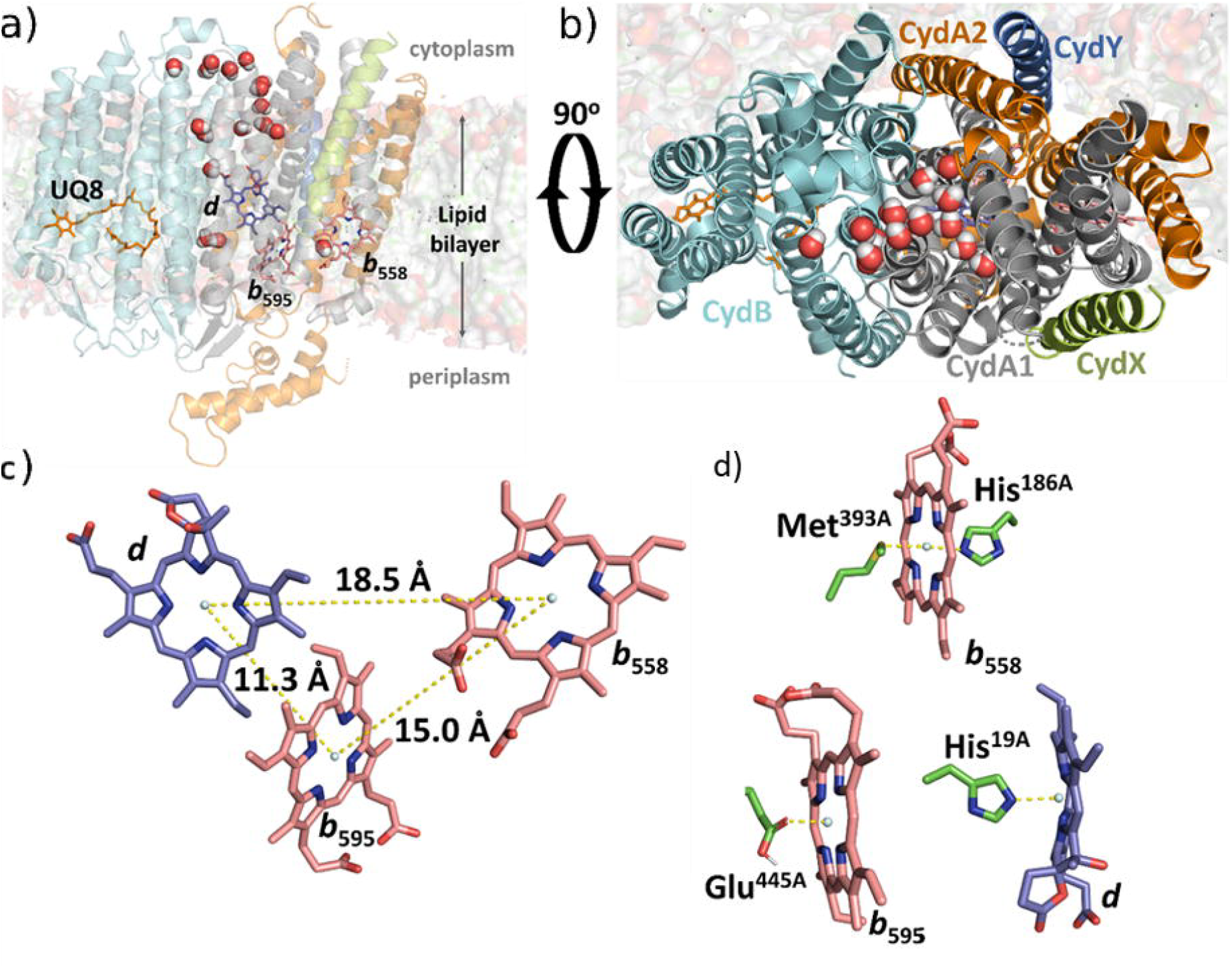
a) computational model of E. coli cytochrome bd inserted in a lipid bilayer, a few water molecules penetrating the protein are shown for illustrative purposes. b) zoom on the protein showing the five protein segments defined in our computational set-up. c) spatial arrangement of the three hemes with Fe-Fe distances determined from the PDB 6RX4 structure. d) illustration of the iron’s coordination spheres without considering here the possibility of water coordination. The protonation states of carboxylic functions are those retained in the present work.

A few previous molecular dynamics simulation or quantum chemistry studies of cytochrome *bd* oxidase have been reported. Safarian *et al*. investigated solvent accessibility in *M. tuberculosis* cyt *bd* suggesting the formation of water strings between the hemes *b* [10]. Furthermore, they identified a menaquinone-9 binding site involving heme *b*_595_ and they conducted two 750 ns MD simulations in the oxidized and reduced states to characterize the binding dynamics of the menaquinone. More recently, Seitz *et al*. used MD simulations of cyt *bd*-I from both *E. coli* and *M. tuberculosis* to identify new potential inhibitor pockets or surface-level binding sites and subsequently screened a collection of compounds through molecular docking in the identified regions [15]. Cao and Liao reported instructive quantum chemistry-based study of some aspects of the reaction mechanism of *E. coli* cyt *bd*-I [16]. They used a so-called cluster approach to describe the active site and the immediate surrounding residues and a static approach (*i.e*. the intrinsic thermal motions are ignored). We note that Glu445 was deprotonated in this work. Within their working hypotheses the authors made a plausible proposal for a complete catalytic cycle for O_2_ reduction at heme *d*.

Many questions, however, remain to be clarified, such as the factors that contribute to an efficient inter-heme electron transfer and the possible electron pathways. The proposed mechanisms also need to be tested using numerical approaches that explicitly consider the heterogeneous nature of these molecular systems and their multiscale dynamics [17]. We make a reasonable assumption that after a sequential 2-electron oxidation of a quinol molecule by heme *b*_558_, the system is left with two electrons on each of hemes *b*. We are thus interested in this work in the subsequent heme *b*_595_ to heme *d* electron transfer that would enable O_2_ binding and its first reduction step. We are also interested in ascertaining the effect of Glu445 protonation on ET thermodynamics. We consider this long range electron transfer to take place within the non-adiabatic, high-temperature limit of the Marcus theory of electron transfer [18–20]. The rate is given by equation 1:

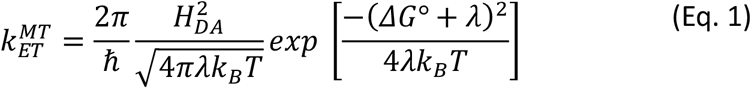

with *ΔG*° and *λ* the free energy of the reaction and the reorganization energy respectively. *H*_*DA*_ is the quantum mechanical coupling matrix elements between the initial and final electronic states. 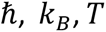 are the reduced Planck constant, Boltzmann constant and the temperature, respectively. In the present work, we evaluate these parameters by means of realistic MD simulations, collecting a total 10 μs of simulation time. We reveal the role played by the protein, by the lipid membrane and other constituent molecules on the determination of free energy and free reorganization energy, in qualitative agreement with the experimental redox potentials. The quantum mechanical coupling is evaluated employing density functional theory calculations and rationalized pathway model analysis [21,22].

## 2. Simulation methods

### 2.1 System preparation

The model system was built with the assistance of CHARMM-GUI (Chemistry at HARvard Macromolecular Mechanics - Graphical User Interface) [23–25] taking the Cryo-EM (6RX4) structure from the Protein Data Bank [5] as a reference. Cytochrome *bd-*I contains two main chains, CydA and CydB, and a small CydX subunit (). The CydA’s structure lacks coordinates for residues 1, 263 to 302 (Q-loop) and 515 to 522, while CydX lacks coordinates for residues 32 to 37. These missing residues are expected to be located in contact with the solvent and the reconstruction of these missing coordinates was not attempted. We denote as CydA1 the CydA’s sequence before the Q-loop (residues 1 to 262) and as CydA2, the sequence after the Q-loop (> 303). On the other hand, the CydY’s coordinates, which are missing in the 6RX4 structure have been borrowed, after structure alignment, from the 6RKO structure [26], albeit with missing coordinates for residues 1 and 2.

Ubiquinone-8 (UQ8) molecule present in the structure was retained in the model (Figure 1). On the other hand, the glycerophospholipid located at the surface of CydB was not retained, because the modelled membrane lipid molecules are assumed to occupy a similar position. The overall structure was oriented according to the “Orientations of Proteins in Membranes convention” thanks to the online server [27] before switching to CHARMM-GUI [25]. The scripts generated via CHARMM-GUI were extracted and rerun on our local cluster with CHARMM version 40b [23] and CHARMM36 forcefield parameters [23]. The reason for this procedure was the need to correctly set the non-standard heme coordination, which was not parameterized by default on the online server. The protonation states of residues Glu74, Glu99, Glu438, Glu489, Glu445, Asp247 from CyA and Asp29, Asp58 from CydB were set according to PropKa calculation [28]. All histidine residues were protonated on δ positions.

Parameters for UQ8 were generated by the CHARMM-GUI server by analogy with other molecules. Heme cofactors were parametrized based on Density Functional Theory calculations (see Supporting Materials for details). Two charge repartitions were investigated as summarized in Table 1. Heme *b*_558_ is always considered to be reduced, while hemes *b*_595_ and *d* are alternatively oxidized or reduced in our simulations. The two states 1 and 2 correspond to the initial and final redox states associated to heme *b*_595_ to heme *d* ET. As illustrated in Table 1, electron transfer involves the displacement of a +1 charge from the latter to the former.

**Table 1:** The two charge repartition schemes considered in this study where R and O stands for Reduced and Oxidized respectively. The numbers in brackets indicate the force field charges on the hemes, without counting here propionate groups’ charges.

| | heme $b_{558}$ | heme $b_{595}$ | heme $d$ |
| --- | --- | --- | --- |
| state 1 | R (-2) | R (0) | O (+1) |
| state 2 | R (-2) | O (+1) | R (0) |

The protein system was inserted into a lipid bilayer containing 75% of PPPE (1-palmitoyl-2-palmitoleoyl-*sn*-glycero-3-phosphoethanolamine), 20% of PVPG (1-palmitoyl-2-vacenoyl-*sn*-glycero-3-phosphoglycerol) and 5 % of cardiolipin (1,3-*bis*(*sn-*3’-phosphatidyl)-*sn*-glycerol) (PVCL2, that is PVPG+PVPG and head group charge of -2). Based on the protein packing, the upper and lower leaflets were made slightly asymmetric, containing 185 PPPE, 48 PVPG and 12 PVCL2 lipids in the upper leaflet and 182 PPPE, 48 PVPG, and 12 PVCL2 lipids in the lower leaflet respectively. The lipids were positioned randomly in each leaflet generating three set-ups, A, B and C with the same membrane composition but different initial random lipid distributions. The membrane protein system was solvated in a hexagonal unit cell with 26 Å width TIP3P [29] water layers above and below the membranes. 306 K^+^ and 163 Cl^-^ counterions were added to ensure electrical neutrality and an ion concentration of 0.15 M. For each set-up A, B and C, the simulations were repeated with different initial velocities. As a result, the individual simulations are labelled A1, A2, …, up to C2 (1 and 2 stand for state 1 and state 2 respectively)

MD simulations were conducted with NAMD (version 2.14) [30]. Each system (A1, A2, …) was equilibrated by a standard computational procedure. This involved a typical heating and equilibration process, where restraints were initially applied to the heavy atoms of the lipids and proteins before being progressively released.

### 2.2 Density Functional Theory

The geometry of the three heme cofactors was optimized in gas phase with Auxiliary Density Functional Theory (ADFT) [31] with the deMon2k program (version 6.1.6 or 6.2.2 [32]). For geometry optimizations we used the TZVP-GGA (Triple Zeta with Valence and Polarization, calibrated for Generalized Gradient Approximation functionals) basis set on all atoms and the GEN-A2* auxiliary basis set [33]. The three molecules were optimized in the gas phase, both in the ferric and ferrous redox states considering three spin states (singlet/triplet/quintet states for Fe^2+^ and doublet/quartet/sextet for Fe^3+^) respectively.

The initial geometries were extracted from the 6RKO structure in the Protein Data Bank[26]. The models retained the amino-acid residues coordinating the iron cation while the C_*α*_ carbon atoms were replaced with hydrogen atoms. In the gas phase, molecular orbitals of the carboxylate groups in the propionates have higher energies than the iron d orbitals making them more likely to be ionized. To address this issue in and enable the calculation of both Fe^2+^ *and* Fe^3+^ states for redox property analysis, the propionates were protonated in the gas phase DFT model. For heme *b*_595_ and heme *d*, geometry optimizations were performed in the presence and absence of a water molecule in an attempt to explore the potential sixth coordination. The optimized structures without water binding were used to derive the force field charges.

Geometry optimizations in the gas phase have been carried out with PBE (Perdew Burke Erzenhof) and single point energy calculations were subsequently done with ORCA [34] software using the *ω*B97X-D3[35–37] XC functional. For the latter, we considered the def2-SVP (C, N, O, H atoms) and def2-tzvp (Fe) basis sets with the def2/j auxiliary basis for resolution-of-the-identity.

### 2.3 Electronic coupling calculations

The electronic coupling *H*_DA_ entering the Marcus rate expression was evaluated by the Projected-Orbital-Diabatization method (POD) as introduced by Thoss and co-workers [38] and more recently popularized by Blumberger and co-workers for applications to hemes mediated electron transfers [39,40]. Briefly, the method can be seen as a post-self-consistent-field method whereby the Hamiltonian matrix is block-diagonalized over the donor and acceptor orbitals to obtain diabatic electron transfer states. The electronic coupling is extracted after transformation of the Hamiltonian matrix in the diabatic basis.

The POD approach was implemented for this work in the deMon2k program (version 6.0.2), ensuring compatibility with the existing QM/MM (Quantum Mechanics/Molecular Mechanics) framework [41,42]. We extracted geometries from the last 400 ns MD trajectories on state 1 every 50 ns, thus generating an ensemble of 48 geometries in total. For each of them, we first conducted 80 steps of geometry optimization to avoid large distortions of the hemes before calculating the POD coupling. We used a TZVP basis-set on all atoms and GEN-A2(C, H)/GEN-A2*(Fe, N, O) auxiliary basis set. The electronic couplings were calculated with the PBE XC functional.

### 2.4 Electron transfer pathway

The pathway model elaborated by Beratan and co-workers [21,43] was used to rationalize electron tunneling based on the molecular structure. The model assumes the electronic coupling to decay along a specific pathway. The overall decay is expressed as product of a constant contact coupling 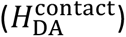 and a factor (*ε*_*tot*_) that accounts for the attenuation of the coupling due to the intervening medium between the two hemes. The factor *ε*_*tot*_ depends on the number of covalent bonds (*N*_c_), hydrogen bonds (*N*_hb_), and through-space jumps (*N*_ts_) that make up the electron transfer pathway, each making a specific contribution to the overall decay (*ε*_c_, *ε*_hb_ and *ε*_ts_ respectively). This factor is computed using a set of mathematical expressions that were empirically calibrated by the model’s authors.

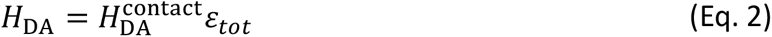

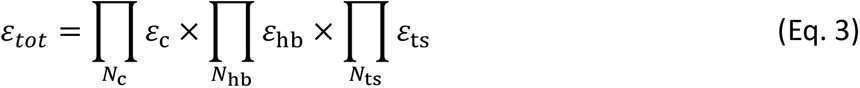

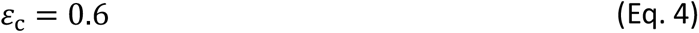

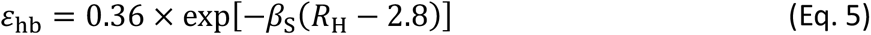

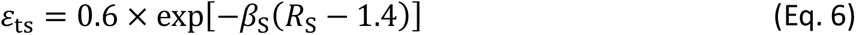

In these equations, *R*_S_ is the atom–atom distance for through-space jumps, *R*_H_ is the hydrogen bond length (taking the inter-heavy atom distance) and *β*_S_ is a characteristic decay factor set to 1.1 Å^-1^ in our computations [22]. The electron transfer pathway analysis [21] was carried out with the in-house “et_paths” module of the Cuby3 framework developed in collaboration with J. Rezac [44]. The plugin identifies the tunneling pathways and calculates the overall decay factor. The iron atoms from heme *b*_595_ and heme *d* were defined as donor/acceptor atoms. The MD trajectories for electronic states 1 and 2 were analyzed, extracting geometries every 20 ns over the last 400 ns of each simulation. The search for the most efficient pathway was carried out with the Dijkstra’s algorithm [45] on pruned geometries.

## 3. Equilibration of cytochrome *bd*-I in explicit membranes

### 3.1 System equilibration and stability assessment

The simulations were run for 780 ns with a 2 fs time step at 303 K and under 1 atm for each redox state and for the two membrane initial configurations. The Root-Mean-Square-Deviation (RMSD) of protein’s backbone atoms and membrane thickness all indicate stability of the molecular systems over the simulations. (Fig. S1-S2). We also computed the RMSD for the heme *b*_595_’s atoms after superimposition on the protein backbone (Fig. S3) to monitor the position of the hemes inside the protein matrix. The value lies between 1 and 5 Å (Fig. S3), which reveals some flexibility. The largest values are associated with slight rotations of the heme within the average porphyrin plane. Heme *b*_595_, perhaps because of its proximity to the surface, has a little space in the protein matrix to accommodate a range of slightly different conformations. Heme *d* is a little less flexible with RMSD values contained below approximately 3 Å, while heme *b*_558_, that is hexacoordinated with Met393 and His186, is the less flexible with a RMSD largely below 3 Å. The structures initially deprived from water molecules readily gets filled by water molecules within roughly 200 ns (Fig. S4). We note some disparity among the different systems in term of protein water content. After 800 ns, for all simulations but C1 in state 1, the protein contains between 45 and 85 water molecules within a cylinder of radius 10 Å and height 20 Å centered on heme *d* iron atom and perpendicular to the membrane.

### 3.2 hemes wettability

The wettability of protein pockets containing hemes was evaluated by counting the number of water molecules in close proximity to the iron atoms. In our set-up, no force field bonding term was added between a water molecule and the iron atoms. Yet, water molecules can come in bonding distance due to the stabilizing electrostatic interaction between the iron atoms and the TIP3P oxygen atoms. For heme *d*, a water molecule is most often found within less than 3 Å from the iron atom (Fig. S5). In MD trajectories corresponding to state 1 (oxidized heme *d*), a water is in close contact to iron atom. Simulation SB1 is an exception. In that case the side chain carboxylate atom from Glu99 interacts with the iron atom instead of a water molecule. For state 2, the likelihood to find a water molecule close to iron is less, but sill substantial (from 30 to 80% of the time).

For heme *b*_595_, we have found that in state 1 (heme reduced), for all simulations, the CydA Phe12 oxygen backbone atom is always in direct interaction with the iron atom and is never displaced by a water molecule within 800 ns. In state 2 (oxidized heme), on the other hand, a water molecule is always found within 3 Å of the iron atom. The force field charges on iron amount to 1.45 and 0.96 in the oxidized and reduced forms, while those on oxygen atoms from Phe12 backbone and TIP3P molecules amount to -0.51 and -0.83 respectively.

## 4. Electron transfer between heme *b*_595_ and heme *d*

### 4.1 Electron transfer theory

The thermodynamics parameters *λ* and *ΔG*° are accessible from the knowledge of the average energy gaps through the Linear Response Approximation (LRA)[46].

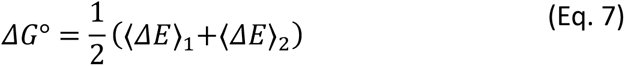

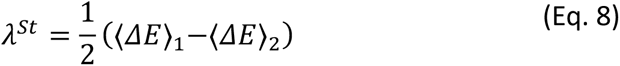

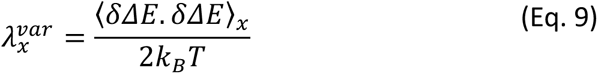

In these equations, ⟨… ⟩_*x*_ denotes an average performed on a sample of structures extracted from a MD simulation carried out on the potential energy surface corresponding to state *x* (1 or 2). The *St* superscript in (Eq. 8) emphasizes the relationship of the reorganization energy calculated with this formula with the Stokes shift encountered in spectroscopy. *ΔE* = *E*_2_ − *E*_1_ is the vertical energy gap between states 1 and 2, that is the potential energy difference between the two states for a fixed given molecular configuration. The potential energy varies because of the electronic structure on the heme cofactor, *i.e*. the change in redox state, and because of the environment (protein, lipid, water …). *δΔE* = *ΔE* − ⟨*ΔE*⟩_*x*_ is the energy gap fluctuation along MD simulation performed on electronic state *x*. Under the LRA, the various ways of evaluating the reorganization energy are equivalent, and under the limit of complete configurational sampling, one should expect 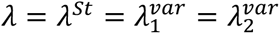. Deviations from this limit equality are quantified by the *χ*_*G*_ parameter. Departures from unit indicate either LRA breaking or incomplete configuration space sampling [47].

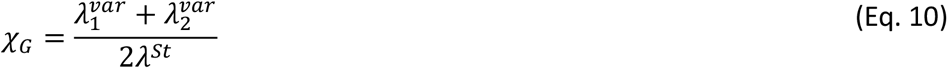

Following previous studies on heme proteins [48–52], we adopt here a QM+MM strategy and decompose the total thermodynamics quantities into inner-sphere and outer-sphere contributions [53]. The former is evaluated on optimized geometries in the gas phase, while the latter is obtained from extensive sampling of system configurations provided by MD simulations. To be more specific, we consider in the inner-sphere of heme *b*_595_ the iron cation, all the atoms of the porphyrin moieties, but the CH_2_COO^-^ propionate groups, and the side chain of Glu445. For heme *d*, the iron cation, all the atoms of the porphyrin moieties, but the CH_2_COO^-^ propionate group, and the side chain His19 are included in the inner sphere. All the interactions between these atoms and the rest of the system is counted in the outer-sphere contribution, in which we also add the inter-heme interactions. It is well-established that classical force fields that do not include explicit electrostatic induction tend to overestimate outer-sphere reorganization energies [53]. On the other hand, when induction is explicitly accounted for, it consistently leads to a reduction of approximately 30%. However, non-polarizable force fields remain faster and allow for more extensive conformational sampling. For this reason, in this study, we have chosen to rely on such force fields and apply a reduction factor of 0.7 to all the outer-sphere reorganization energies reported below.

### 4.2 Electron transfer driving force

Figure 2, left, depicts an example of outer-sphere vertical energy gap fluctuations for both redox states (see Fig. S7 for all the other simulations) while Figure 2, right shows the Marcus parabola obtained by the application of the LRA to the vertical energy gaps (considering only the outer-sphere contribution). After an initial phase of a few hundred of nanoseconds, the *ΔE* values achieve rather steady fluctuations on the timescales of the simulations. The last 400 ns were thus considered for calculating Marcus thermodynamics parameters in our setups.

**Figure 2:**
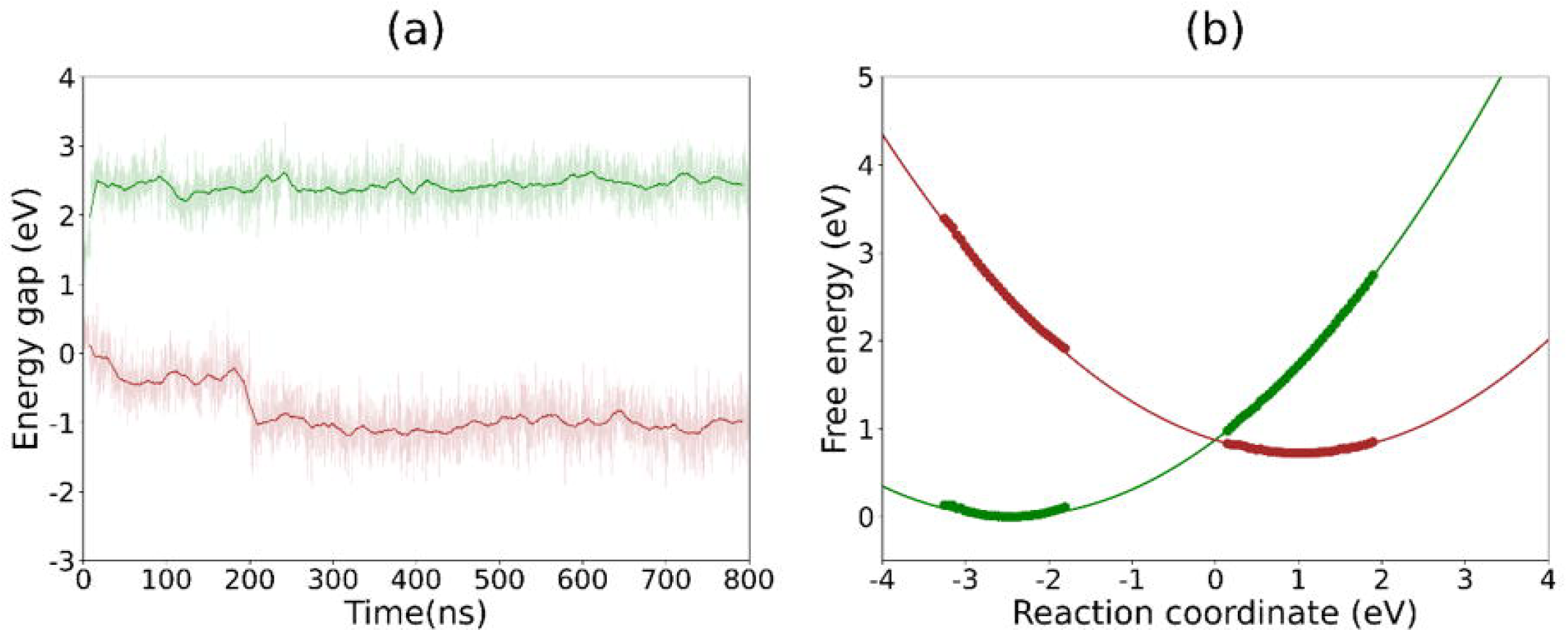
outer-sphere contribution for heme b_595_ to heme d electron transfer for the SA1 set of simulations. a) fluctuations of outer-sphere vertical energy gap. The thick lines are rolling average over 16 ns time windows. b) Marcus parabolas obtained by application of the LRA and ergodic hypothesis for this MD simulation set [46].

Table 2 collects the parameters for all the six sets of simulations. The values of outer-sphere *ΔG*^*o*^ are mostly positive but span a large array, ranging from -0.05 ± 0.07 eV (set SB1) to 0.72 ± 0.02 eV (set SA1) with an average value of 0.34 eV. As *ΔG*^*o*^ is calculated from an additive force field, further insights can be obtained by decomposing the overall free energy over the system’s constituents (Figure 3, top and Fig. S8). This decomposition shows that the protein scaffold itself either favor or disfavor ET depending on the simulations sets. While CydB and CydH favor electron transfer in all simulation sets, CydA2 (i.e. CydA after the Q-loop, residues 303 to 515 in our set-up) strongly disfavor ET. These trends can be rationalized considering the atomic charges of the redox cofactors (Table 1).

**Table 2:** Outers-sphere and inner sphere thermodynamic parameters calculated with ωB97XD functional. The numbers in brackets correspond to block-average uncertainties calculated over five blocks. * a “w” sign before the heme indicate that we consider a hydrated form. s, t, q respectively indicate a complex in the singlet, triplet or quartet state.

| Setup | $\Delta G^0$ (eV) | $\lambda^{St}$ (eV) | $\lambda_1^{var}$ (eV) | $\lambda_2^{var}$ (eV) | $\chi_G$ |
| --- | --- | --- | --- | --- | --- |
| outer-sphere |  |  |  |  |  |
| SA1 | 0.72 ($\pm 0.02$) | 1.22 ($\pm 0.01$) | 0.69 ($\pm 0.02$) | 1.03 ($\pm 0.05$) | 0.71 |
| SA2 | 0.38 ($\pm 0.05$) | 1.05 ($\pm 0.03$) | 0.80 ($\pm 0.02$) | 1.16 ($\pm 0.09$) | 0.71 |
| SB1 | -0.05 ($\pm 0.07$) | 1.02 ($\pm 0.03$) | 1.09 ($\pm 0.06$) | 1.19 ($\pm 0.05$) | 1.12 |
| SB2 | 0.39 ($\pm 0.02$) | 1.00 ($\pm 0.01$) | 0.88 ($\pm 0.06$) | 1.10 ($\pm 0.08$) | 0.99 |
| SC1 | 0.22 ($\pm 0.03$) | 1.03 ($\pm 0.02$) | 0.79 ($\pm 0.05$) | 1.14 ($\pm 0.07$) | 0.94 |
| SC2 | 0.36 ($\pm 0.02$) | 1.33 ($\pm 0.01$) | 0.75 ($\pm 0.03$) | 0.99 ($\pm 0.04$) | 0.65 |
| inner-sphere* | $\Delta E^{is}$ | $\lambda^{is}$ | | | |
| $b_{595}^{II,t} + d^{III,q} \rightarrow b_{595}^{III,q} + d^{II,t}$ | 0.27 | 0.28 | - | - | - |
| $b_{595}^{II,t} + wd^{III,d} \rightarrow b_{595}^{III,q} + wd^{II,s}$ | 0.13 | 0.22 | - | - | - |
| $wb_{595}^{II,t} + d^{III,q} \rightarrow wb_{595}^{III,q} + d^{II,t}$ | -0.18 | 0.60 | - | - | - |
| $wb_{595}^{II,t} + wd^{III,d} \rightarrow wb_{595}^{III,q} + wd^{II,s}$ | -0.32 | 0.67 | - | - | - |

**Figure 3:**
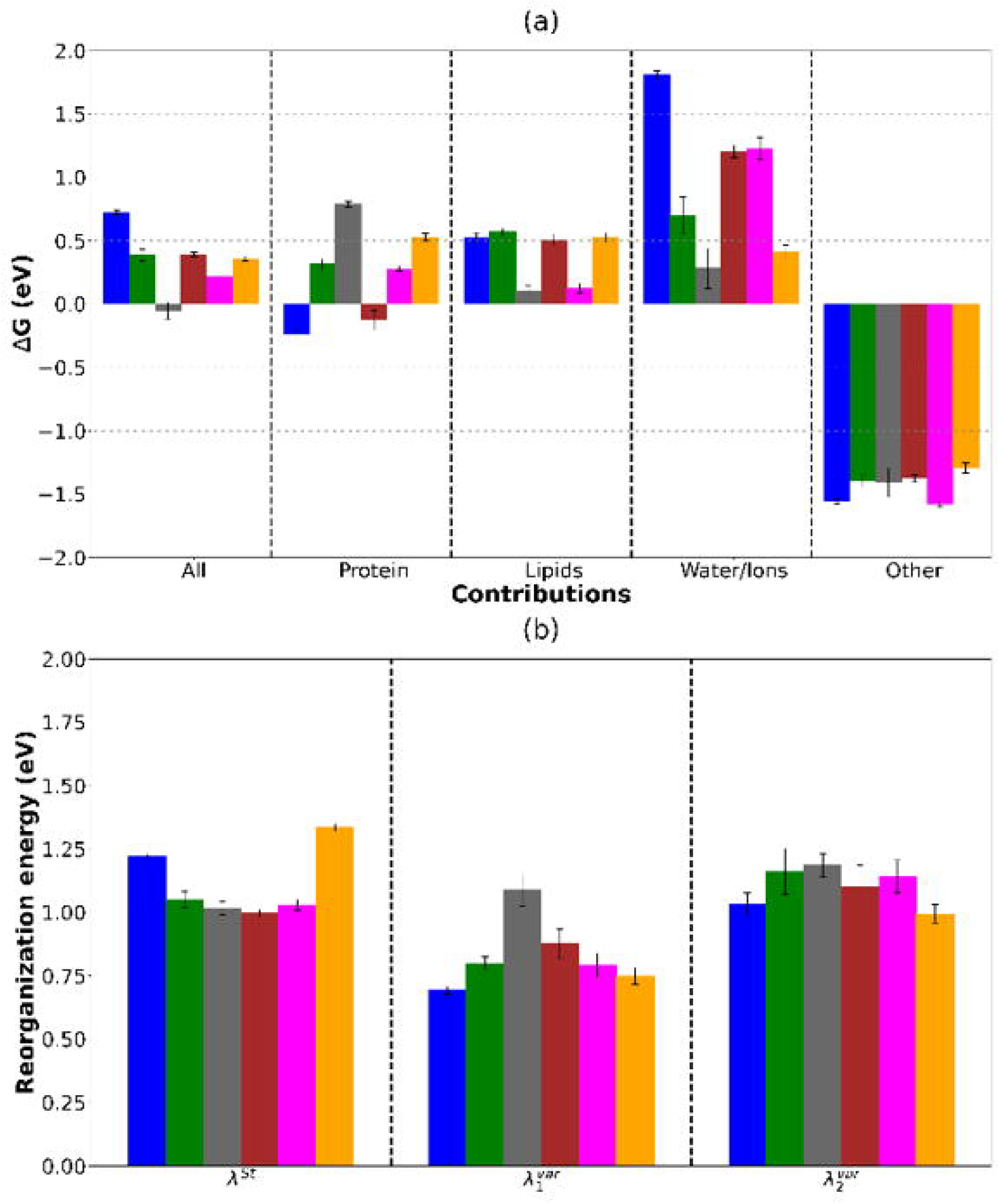
a) decomposition of ΔG^°^ over the constituents of the system, namely the protein, lipid membrane, water and counterions and cofactors (i.e. heme b_558_, propionate groups, UQ8 and inter-heme interaction). b) outer-sphere reorganization free energies as computed from (Eq. 8) (first block), and (Eq. 9) in the 2 redox states (second and third blocks). In each block the six vertical bars correspond to sets SA1 (blue), SA2 (green), SB1 (grey), SB2 (brown), SC1 (magenta), and SC2 (orange).

Examining the contributions from the lipid membrane, we note that the upper and lower leaflet show similar, disfavoring effect, on the driving force (0.15-0.5 eV). In fact, the initial random partitioning of lipids in the leaflet (sets A, B and C) has less impact than the initial conditions used for a given set (i.e. comparing B1 vs B2 or C1 vs. C2). Further investigations revealed that the most abundant (75%) PPPE lipids holding no net electric charge from the upper or lower leaflet have similar and slightly disfavoring contributions (Fig. S8). On the other hand, the PVPG and cardiolipin (PVCL2) molecules respectively holding net charges of -1 and -2 per molecule, have greater, but opposite effect on the driving force depending on their positioning in the upper (unfavorable effect) or lower (favorable effect) leaflets. Inter-heme ET results in +1 charge displacement from the heme *d* to heme *b*_558_. As heme *d* is buried deeper into the protein than heme *b*_595_ (Table 1), the electrostatic interaction with the negatively charged lower leaflet is more stabilizing, hence produces a negative contribution to *ΔG*^*o*^. The same reasoning applies, but in opposite way for the upper leaflet.

Similarly, heme *b*_558_ that holds a negative -2 charge (including that of the propionate groups) logically favors ET (on average -0.57 eV – data given in Table S10 of Supplementary Information) as a consequence of the +1 charge lying on heme *b*_595_ after ET and shorter inter-heme *b* distances (Figure 1). The contribution from heme *b*_595_ and heme *d* propionates is strongly favorable (-0.86 eV in average – data given in Table S9 of Supplementary Information), which has to be understood from the fact that hemes *b* hold two propionate groups (negatively charged in our set-up) while heme *d* contains only one propionate and a *γ*-spiro-lactone ring. Moreover, the relative positioning of the propionate groups favors electrostatic interaction in final electronic state.

Water molecules and counter-ions disfavor ET (Figure 3), but a closer look at the decomposition (Fig. S7 bottom-left) shows that most of the effects comes from the potassium counterions presumably lying close to the lower leaflet. As discussed above the lower leaflet favor ET, but this effect is tempered by the presence of K^+^ counterions that disfavor the transfer. These compensatory effects of thermodynamic contributions between the membrane and the solvent (water and counterions) have also been recently demonstrated by simulations of NADPH oxidases [54].

### 4.2 Reorganization free energies

The reorganization energies are illustrated in Table 2 and Figure 3 (bottom). The outer-sphere contribution ranges from 1.00 to 1.33 eV using Eq. 8, and from 0.69 to 1.19 eV using Eq. 9 (taking into account the 0.7 reduction factor as mentioned in the Simulation methods section). The ergodic parameters are comprised between 0.65 and 1.12, which indicates that the conformational space is not fully explored by our simulations despite a total duration of almost 10 *μ*s in total. This highlights the challenges of reaching extensive conformational sampling of complex molecular systems such as the one investigated here. Electron transfer modelling cannot thus rely on a unique set of MD simulation, but requires replicating the simulations with different initial conditions to obtain sufficiently robust trends. We find reorganization energies of 1.1 eV on average, which is rather typical of what one would expect for a biological inter-heme ET. Looking more closely at the decomposition (Table S3), we find that the protein is the main contributor to *λ* ^*St*^, and in particular the CydA chain (Table S5). The degree of reorganization of water and counter ions is variable, ranging from 0.1 eV for SA2 and SB2 to 0.8 eV for SA1. On the other hand, the membrane contributes less to reorganization energy (< 10%). Reorganization due to other cofactors including heme *b*_558_, propionate groups, UQ8 or inter-heme are gives positive or sometimes negative values depending on the simulation set. Negative values of Stokes reorganization energy indicate that the dynamics of these sets of atoms are not solely modulated by the redox state change. In fact, for small groups of atoms, the lambda decomposition has limited relevance.

### 4.3 Inner-sphere contributions

We now move to the inner sphere contributions as calculated by DFT. We have considered first heme cofactors with or without water coordination. The inner-sphere contributions are calculated at the DFT level following ref. [53]. *λ*^*is*^ has been calculated as:

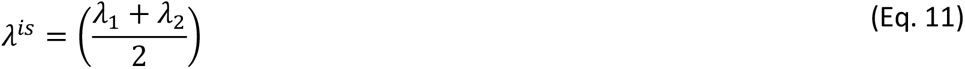

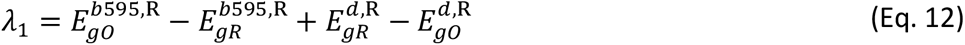

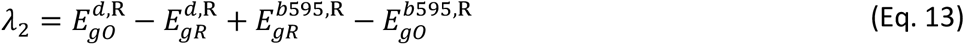

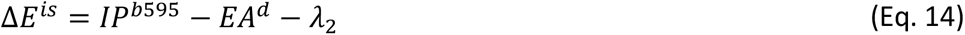

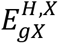 stands for the energy of heme *H* in its redox state *X*(*O or R*) and at the equilibrium geometry *gX*^′^ corresponding to redox state *X*^′^. *IP*^*H*^ and *EA*^*H*^ are the ionization potential and electron affinity of heme *H*, respectively. All the results are collected in Table S1.

The five coordinated heme *b*_595_ is found to be high spin in both the reduced and oxidized states (triplet and quartet, respectively) with ωB97X-D3 XC functional. In the ferrous state, water coordination brings the three spin states close in energy. A point of caution has to be raised here about the difficulty of DFT to correctly predict spin states splitting in hemes. In the reduced, triplet state, water is not bound to the iron (d(Fe-OH_2_) = 3.31 Å), nor is it the case in the oxidized quartet state, or with loose binding d(Fe-OH_2_ = 2.44 Å). On the other hand, water is tightly coordinated in low-spin states (singlet and doublet with d(Fe-OH_2_ of 2.04 and 1.99 Å) respectively. As a consequence, the inner-sphere reorganization energies are large (>0.8 eV) due to water binding/unbinding.

The results for heme *d* follow those for *b*_595_. In absence of water in the coordination sphere, the intermediate triplet and quartet spin states are found to be the most stable in the ferrous and ferric stats respectively. Coordination of water seems possible in the reduced, singlet state.

Considering water free complexes, we find that the respective ionization potential and electron affinity give rise to a positive inner-sphere contribution (Δ*E*^*is*^) of +0.27 eV and a reorganization energy of 0.28 eV (*λ*^*is*^), see Table 2 and Table S1. These values can be directly summed to the outer-sphere contribution and indicate an unfavorable transfer. On the other hand, as we have optimized heme structures including also water molecules in the coordination sphere, we can assess the effect of such coordination on the inner sphere contribution. Considering water binding to heme *d* decreases Δ*E*^*is*^ to 0.13 eV, which is due to the fact that heme *d* is now low spin in the most stable spin configuration (singlet and doublet). Now, considering water coordination to heme *b*_595_, a further drop to -0.18 eV is obtained for Δ*E*^*is*^ and to -0.32 eV when each heme is hydrated. This comes at the price of a large inner-sphere reorganization energy larger than 0.60 eV that can be traced back to the coordination of water itself to the iron cation of heme *b*_595_ (see above). Indeed, in the ferrous, triplet state, water is unbound and has to come to bonding distance in the ferric state. As seen in our MD simulations, water molecules are present in the vicinity of the iron cation making this scenario a likely possibility (Fig. S5). The combination of inner-sphere and outer-sphere contributions in the case of hydrated hemes is not straightforward as the latter already partially account for electrostatic effect of a water molecule close to the iron cations. A simple sum of the two contribution would be plagued by double some counting effect. That said, our DFT calculations reveal the great sensitivity of the inner-sphere contribution to the coordination-sphere contribution and spin state ordering. This in turn suggests possible coupling of inter-heme electron transfer with water coordination associated to spin state reordering.

### 4.4. Quantum mechanical coupling

The electronic coupling has been calculated with the quantum mechanical POD approach at the QM/MM level to consider environment effects (Table 3). We obtain an average value of 0.55 meV with a standard deviation of 0.56 meV, placing the inter-heme electron transfer into the non-adiabatic regime. A pathway analysis gives an average decay ⟨*ε*_*tot*_⟩ of the order of 10^-4^. As depicted in Figure 4, tunneling is dominated by direct heme-to-heme interactions with pathways running alternatively through vinyl and methyl peripheral groups.

**Table 3:**
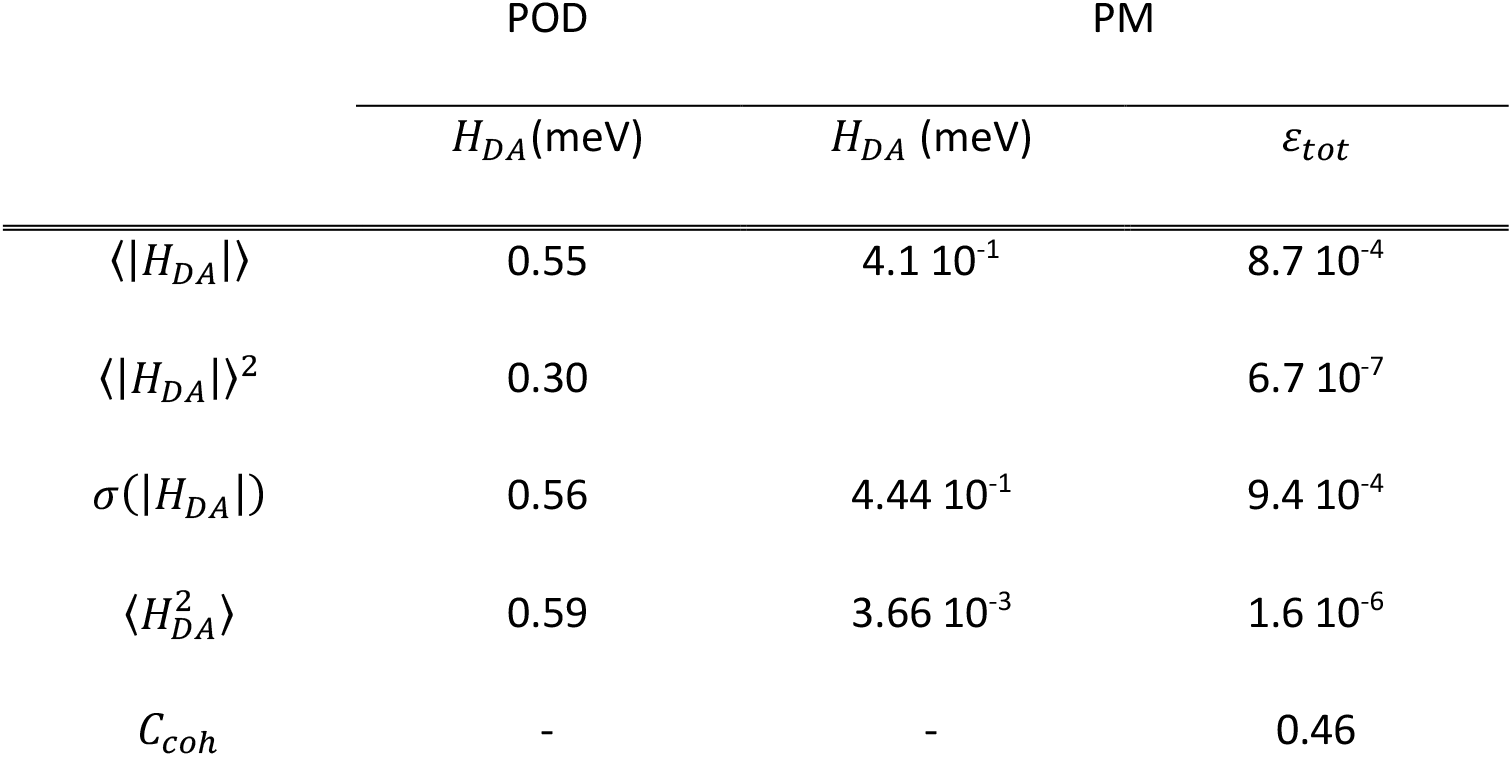
electronic coupling for heme-to-heme tunneling calculated either quantum mechanically with the projector-operator-diabatization (POD) approach or by the semi-empirical pathway model. For the latter a 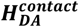 value 47 meV is used [22].

**Figure 4:**
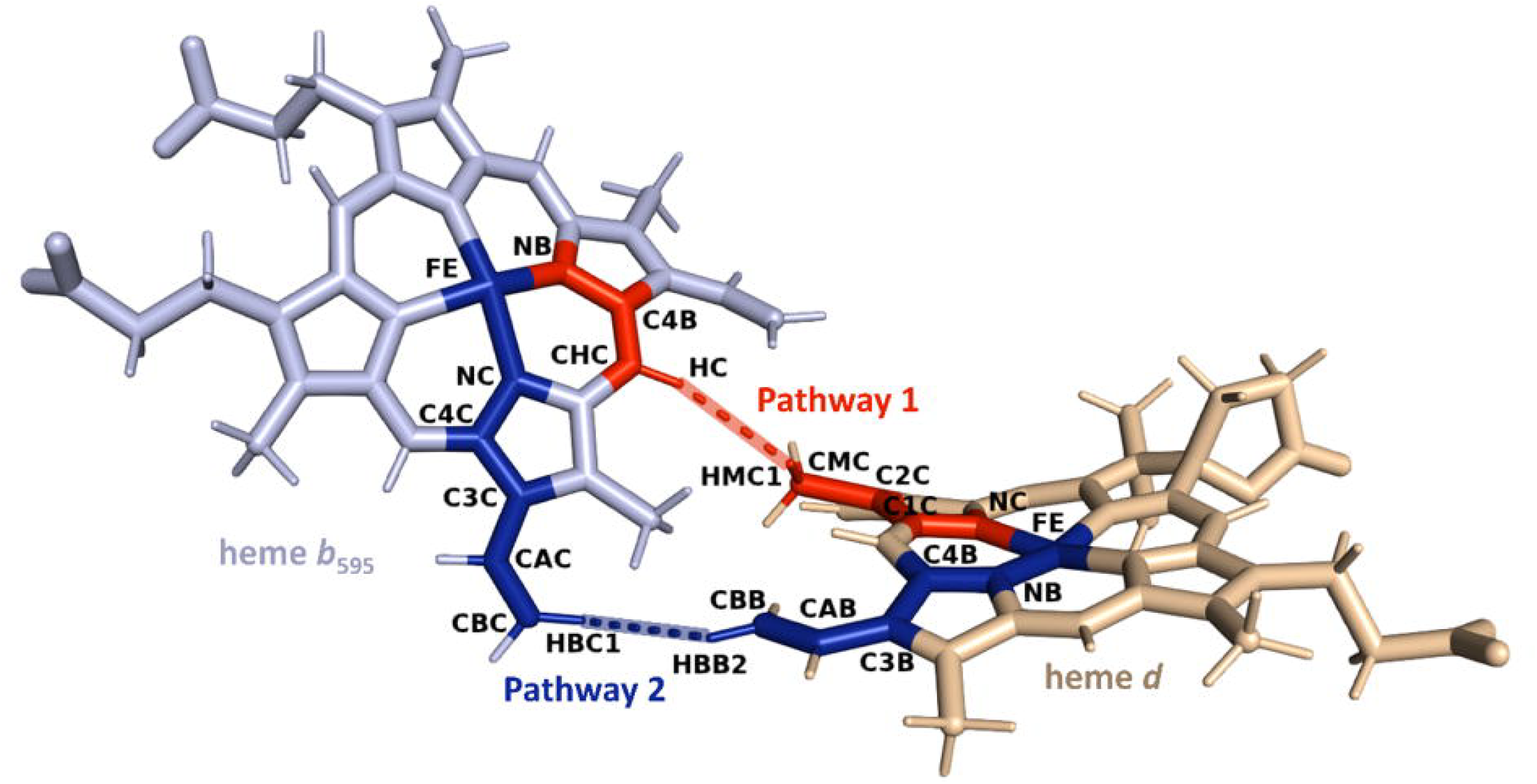
representative electron transfer pathways identified along the MD simulations.

## Discussion and conclusion

The molecular mechanisms by which cytochromes *bd* enable bacterial respiration through their three hemes remain only partially understood, despite significant progress over the past decades, particularly with the publication of atomistic structures from various organisms [4– 10]. The distinct chemical structures of the two *b*-type hemes and the unusual coordination sphere of their iron cations raise important questions, especially concerning the regulatory mechanisms of inter-heme electron transfer. Computational studies can shed light on these processes [20], but to date few have attempted to address these questions for this family of enzymes [10,15,16]. Cao and Liao reported quantum chemistry calculations of the heme active sites relying on the cluster approach [16]. They proposed a sequence of chemical steps involving various proton-coupled electron transfers (PCET). The study provides a plausible catalytic cycle for O_2_ reduction in terms of energetics. Among other conclusions, they suggested that hemes *d* and *b*_595_ could be respectively reduced and oxidized before O_2_ binding. Then, a PCET would take place, to i) reduce *b*_595_, presumably by heme *b*_558_, and ii) to protonate Glu107 by proton transfer from the heme *d* peripheric hydroxyl group. The doubly reduced heme pair (*d* and *b*_595_) would then allow O_2_ binding and reduction. The results reported by Cao and Liao, though valuable and interesting, must be analyzed in light of the assumptions made by the authors to build their model, that is a static cluster approach restricted to heme *d* and heme *b*_595_ cofactors and a few surrounding key residues, and more importantly the absence of dynamic description of the system and its heterogenous composition.

Here, we report a combination of molecular dynamics, quantum mechanics, and hybrid QM/MM simulations of cytochrome *bd* in realistic membranes. This work represents the first computational investigation of electron transfer between the terminal hemes in *E. coli* cytochrome *bd*-I. Our primary objective was to capture the heterogeneous environment of this membrane redox protein. The thermodynamics of electron transfer within a membrane redox protein results from fluctuations in electric fields caused, on the one hand, by the protein scaffold, which is relatively constrained by its primary sequence, by the lipid bilayer, which is inhomogeneous and partially charged, but endowed with a certain fluidity, and by the aqueous and ionic environment, which is highly polar and very flexible. Our objective here was to highlight the role of these different contributors to the driving force and reorganization energy of ET. In addition, we aimed to evaluate the effect of protonating Glu445, the apical ligand of heme *b*_595_, on electron transfer kinetics. We have shown that the overall thermodynamics balance results from large and opposite effects. The CydA domain tend to disfavor ET while CydB and to a lesser extent CydH favor electron transfer, showing that the protein matrix cannot be considered a homogenous continuum for dealing with the inter-heme electron transfer within these proteins. Furthermore, our set-up includes a realistic description of the lipid bilayers, that also is shown to have non-homogenous effect on electron transfer thermodynamics.

We found the outer-sphere contribution for ET between heme *b*_595_ and heme *d* to be unfavorable by a few tenths of eV. Besides the protein matrices, lipids and solvent, we also found contributions from the hemes’ propionate and from heme *b*_558_ to be particularly prominent in driving favorably an electron toward heme *d*. In our simulation set-up, heme *b*_558_ was considered as reduced, hence favoring the transfer of one electron from heme *b*_595_ to the remote heme *d*. Regarding the propionates, the chemical asymmetry of hemes *b* and *d*, favor the reduction of heme *d* when propionates are deprotonated, as in our-set-up. We note that nor the propionate groups, nor heme *b*_558_ were included in the model of Ref. [16]. These major differences prevent a direct comparison of our results with this previous study. On the other hand, it has been suggested based on a cryo-EM structure and MD simulations of *M. tuberculosis* that a solvent filled region near the hemes could constitute a dielectric well that could facilitate deprotonation/protonation of propionates depending on the heme redox states [10]. Our results on *E. coli* cytochrome *bd-*I fully support this view as protonation of one or more of propionates would strongly alter their contributions to the outer-sphere thermodynamics balance. To go deeper, new simulations would be needed to evaluate the propionates’ p*K*_a_ as a function of the heme redox states.

The inner-sphere contribution was calculated considering either five-coordinated hemes or six-coordinated hemes with a water molecule completing coordination. In the first scenario, when considering ET from the doublet heme *b*_595_ to triplet heme *d*, we found unfavorable energetics, and as discussed above, not compensated by the outer-sphere contribution. For an overall charge of 0 and +1 on heme *b*_595_ and heme *d* (without propionates’ charges in this formal counting), the outer-contribution is unfavorable. This suggests that a key factor to favor electron transfer toward heme *d* is either to stabilize reduced heme *d*, maybe by O_2_ coordination and partial reduction toward an Fe(III)/O_2_^•-^ moieties. We also found that water coordination to heme *d* would help to lower the energy balance to 0.13 eV by modifying spin state ordering. Alternatively, one could destabilize ferrous heme *b*_595_, for instance by deprotonating Glu445, which would favor the ferric state. In our MD simulations, as in those of ref. [10] of *M. tuberculosis* cytochrome *bd*, Glu445 (or Glu398 in of *M. tuberculosis*) was protonated. In case of a glutamate form, the outer-sphere contribution would need revision as the protein dynamics would likely be altered by the introduction of the negative charge at heme *b*_595_. Our calculation also showed that water binding to heme *b*_595_ in the ferric state would strongly favor electron transfer. Such a scenario assumes that a water molecule attaches itself to the ferric cation, even though it is not initially bound, leading in term to a large inner-sphere reorganization. We also note that a large reorganization of the coordination sphere is likely to put the ET regime beyond the standard Marcus theory description of electron transfers [19,55].

To summarize, this study provides new insights into the molecular mechanisms underlying the complex redox activity of cytochromes *bd*. Our results suggest a tight coupling between electron transfer and the catalytic steps occurring at the hemes. The composition of the hemes’ coordination spheres—including water, dioxygen, or other ligands—as well as the protonation states of propionates and other key factors, warrant further investigation to fully elucidate the reaction mechanisms at play.

## Supporting information

<Insert Figure 1>

## Acknowledgments

We are grateful to the French *Grand Équipement National de Calcul Intensif* for providing us with generous computational ressources (project numbers A0160706913 and A0140706913).

This work was supported by the Interdisciplinary Thematic Institute SysChem, via the IdEx Unistra (ANR-10-IDEX-0002) and by the CSC Graduate School (CSC-IGS ANR-17-EURE-0016) within the French Investments for the Future Program to RS and PH. PH also acknowledges the Institut Universitaire de France and the Foundation Jean-Marie Lehn.

## Supplementary Information

Graphs monitoring MD simulation stability. Statistics about protein water content. outer-sphere energy gap fluctuations and Marcus parabolas. Tables and Figures displaying contributions to outer-sphere free energy and free reorganization energy. DFT results for the heme cofactor. Python scripts to calculate Marcus theory parameters from vertical energy gaps and Orca software input files to compute inner-sphere contributions can be downloaded on Zotero (10.5281/zenodo.17043776).

