## Supplementary material for "Computational study of heme *b*_595_ to heme *d* electron transfer in E. coli cytochrome *bd*-I oxidase": <Insert Figure 1>

a : UMR 7140, Chimie de la matière complexe, Université de Strasbourg, CNRS, F 67000 Strasbourg

b : Institut de Chimie Physique, Université Paris Saclay, CNRS, F-91405 Orsay, France

### SUPPLEMENTARY INFORMATION

[1] present address : CNRS, Laboratoire de Chimie Théorique, LCT, Sorbonne Université, Paris, France.
[2] present address :

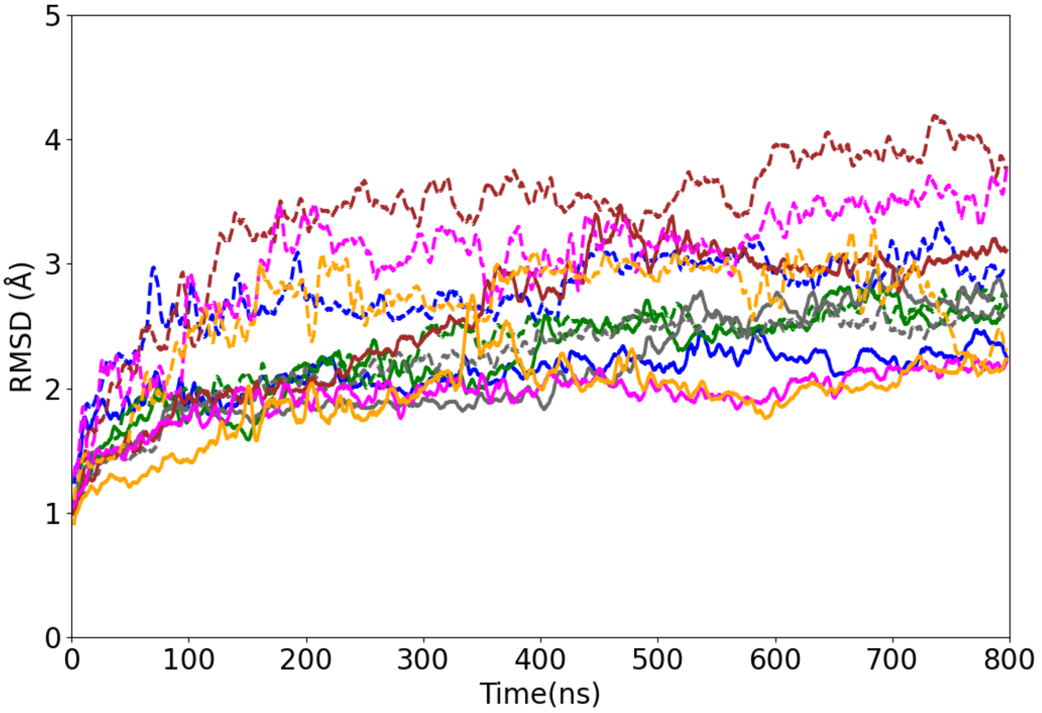


*Figure S1: RMSD of the backbone atoms of cytochrome bd (rolling average over 0.8ns time window). Setup SA1: blue, setup SA2: green, setup SB1: grey, setup SB2: brown, setup SC1: magenta, setup SC2: orange. The plain and dashed curves correspond to the redox states 1 and 2 respectively.*

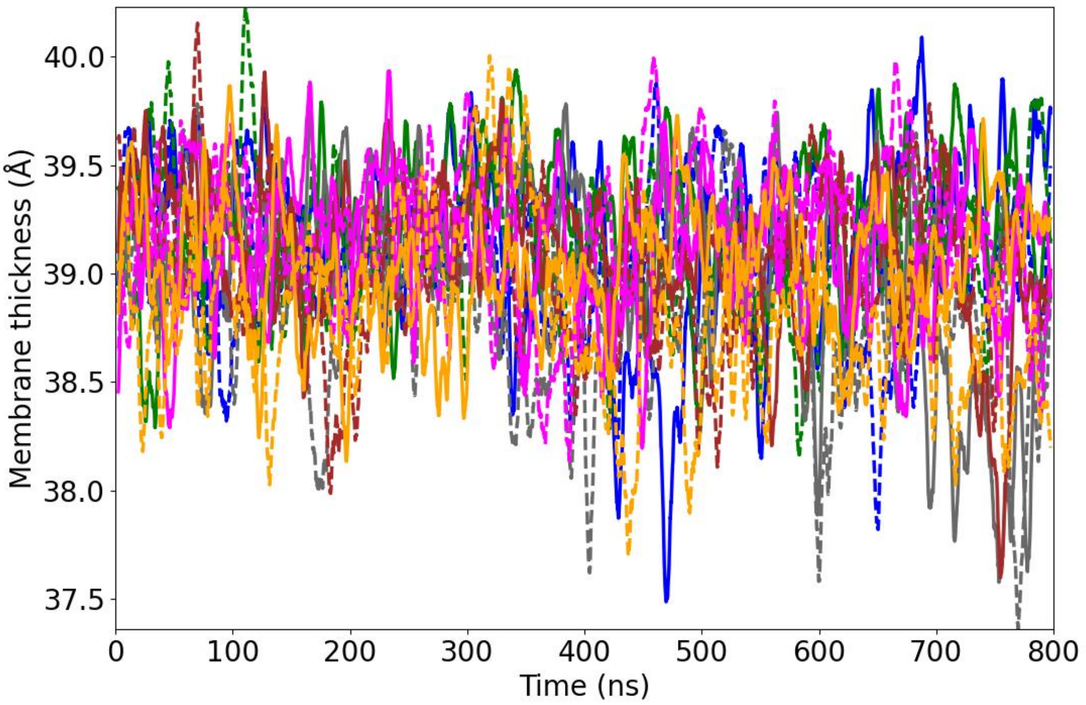


*Figure S2: Membrane thickness, measured as the average distance between the phosphorus atoms of the upper and the lower lipid leaflets. Color code and line types are the same as in figure S1. Setup SA1: blue, setup SA2: green, setup SB1: grey, setup SB2: brown, setup SC1: magenta, setup SC2: orange. The plain and dashed curves correspond to the redox states 1 and 2 respectively.*

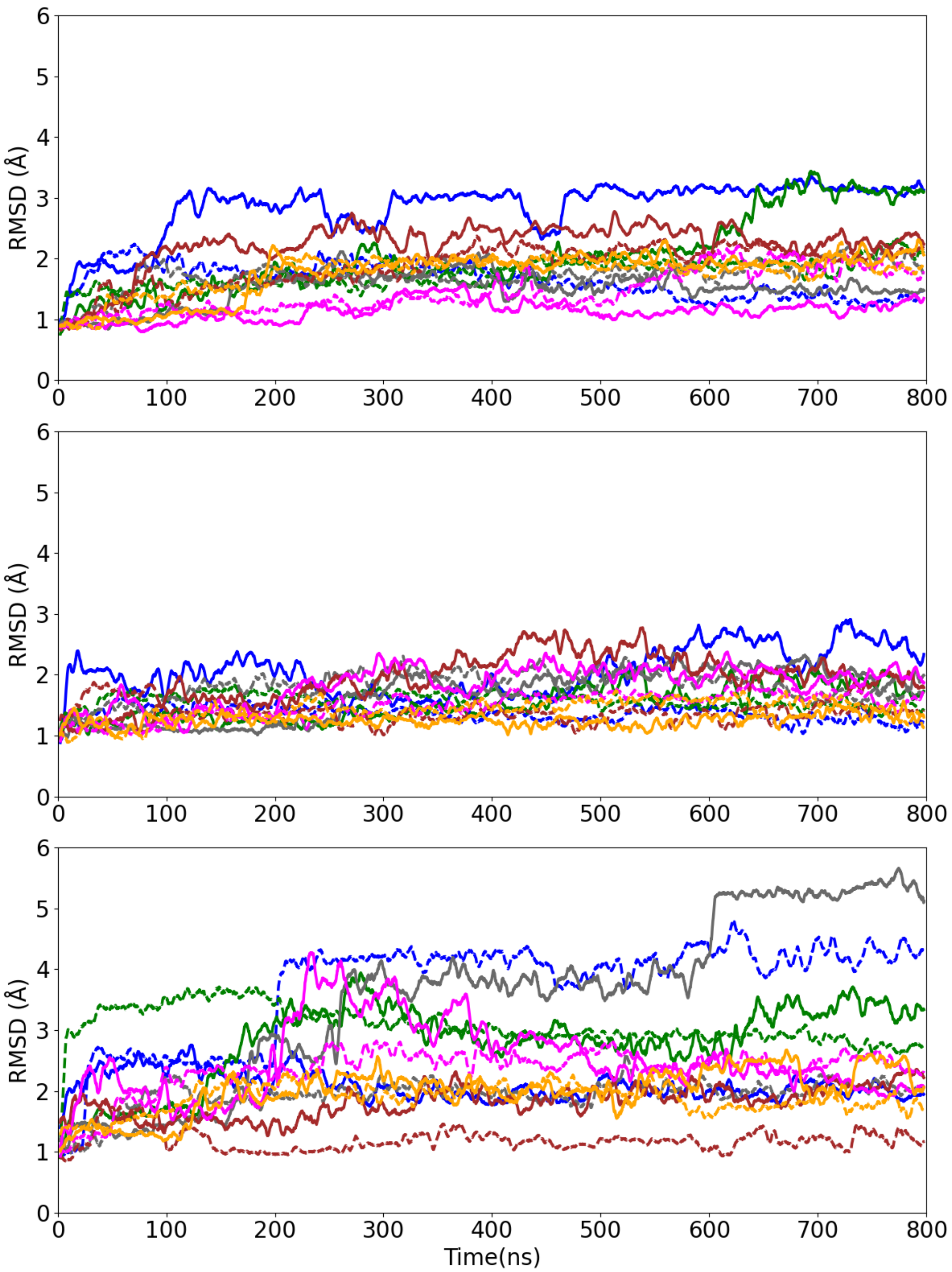


*Figure S3: Structural variability of the hemes (top: heme d, middle: heme $b_{558}$, bottom: heme $b_{595}$). The RMSD of the heme atoms are calculated after a superposition on the protein backbone). Setup SA1: blue, setup SA2: green, setup SB1: grey, setup SB2: brown, setup SC1: magenta, setup SC2: orange. The plain and dashed curves correspond to the redox states 1 and 2 respectively.*

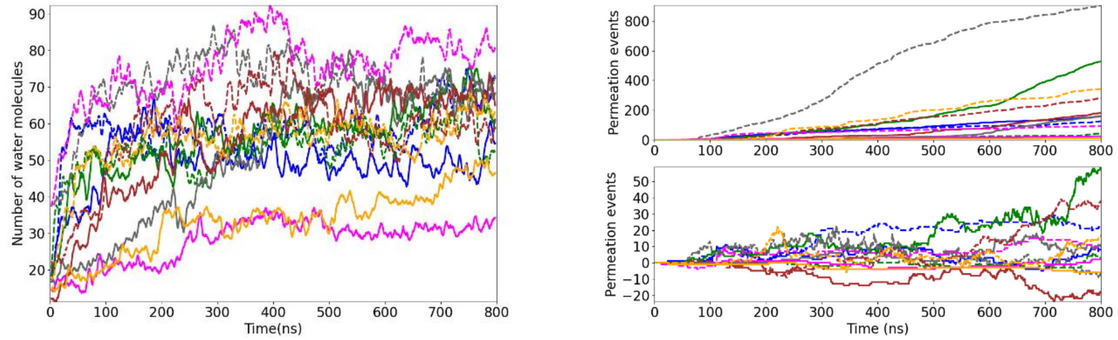


*Figure S4: Number of water molecules in the transmembrane domain, lying in a cylinder of radius 10A and height 20A, centered on Fe of heme D, perpendicular to the membrane (aligned on z axis); Right: permeation events through the protein during the simulation. Upper panel: total number, lower panel: difference between permeation towards cytoplasmic side and towards extracellular side. Setup SA1: blue, setup SA2: green, setup SB1: grey, setup SB2: brown, setup SC1: magenta, setup SC2: orange. The plain and dashed curves correspond to the redox states 1 and 2 respectively.*

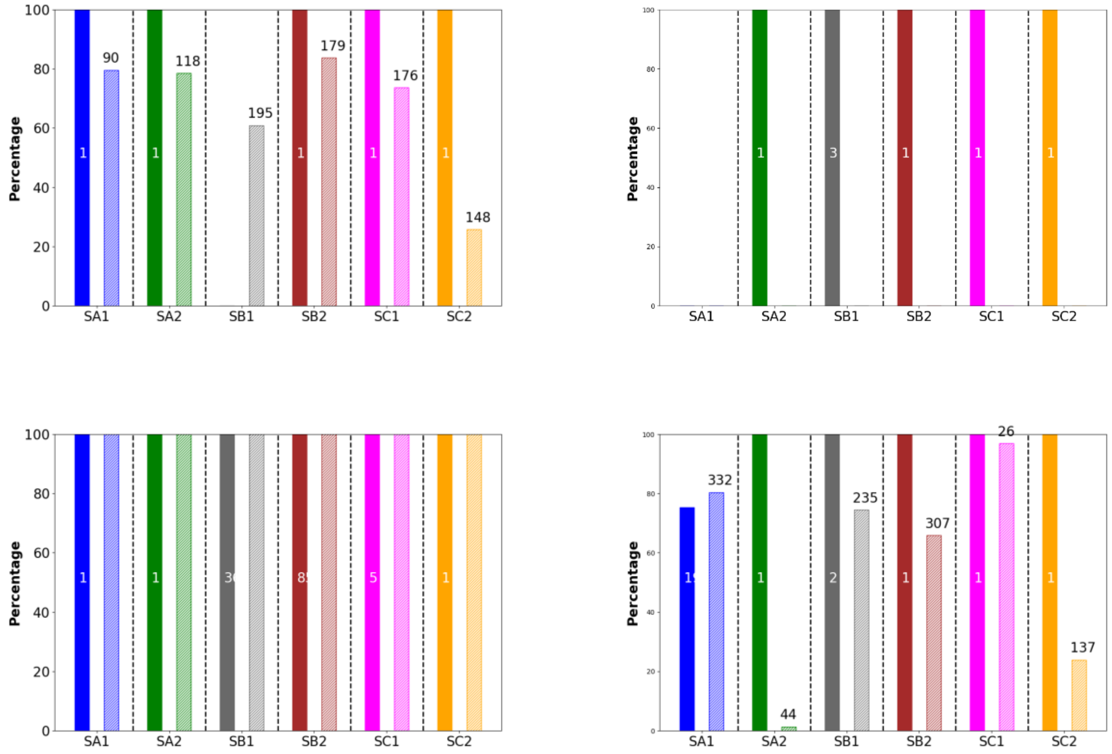


*Figure S5: Statistics of coordination of water molecules to the hemes. Left : coordination to heme d ; Right : coordination to heme b$_{595}$. A water is considered to be coordinated if the distance between the water oxygen and the iron atom is less than 3 Å (upper pannel) or 5 Å (lower pannel). Initial redox state: plain bar ; final redox state: hatched bar. The number given within (initial state) or above the bar (final state) is the number of different water molecules that coordinate the heme.*

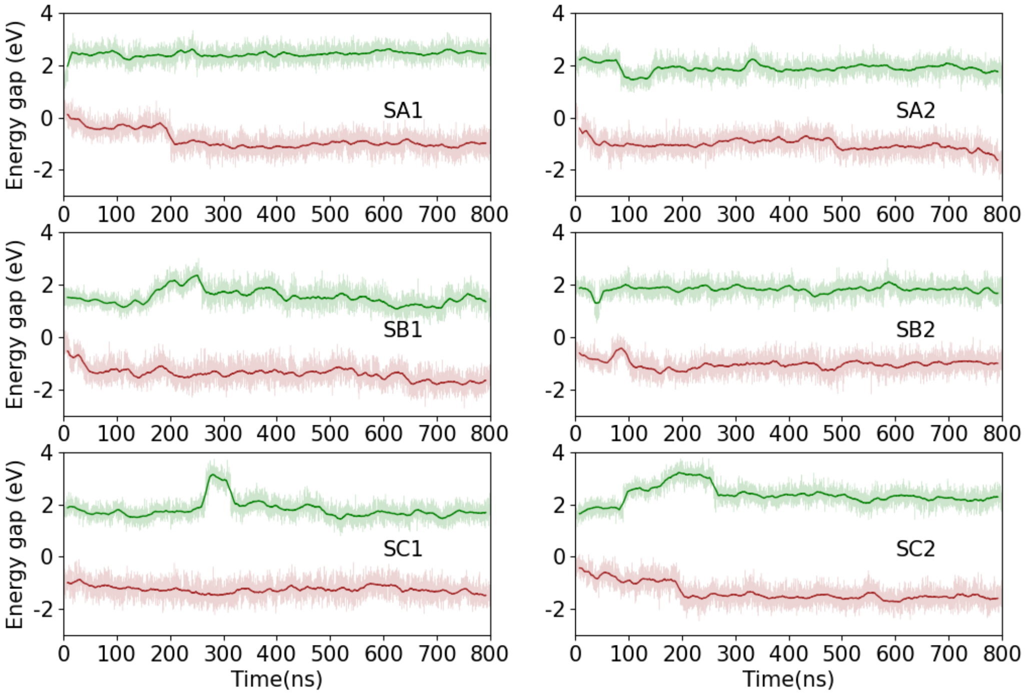


*Figure S6: outer-sphere energy gap fluctuations in states 1 and 8 for each of the 6 MD trajectories. The thick lines correspond to running averages over time windows of 16ns*

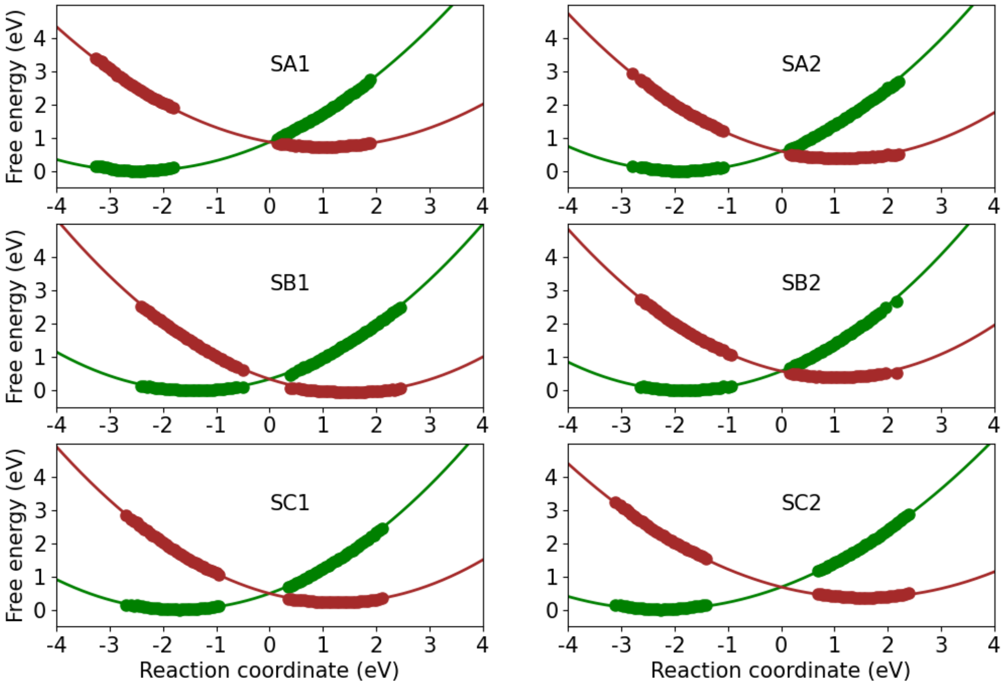


*Figure S7: Marcus parabolas obtained by application of the LRA and ergodic hypothesis to the outer-sphere energy gap for all simulation setups.*

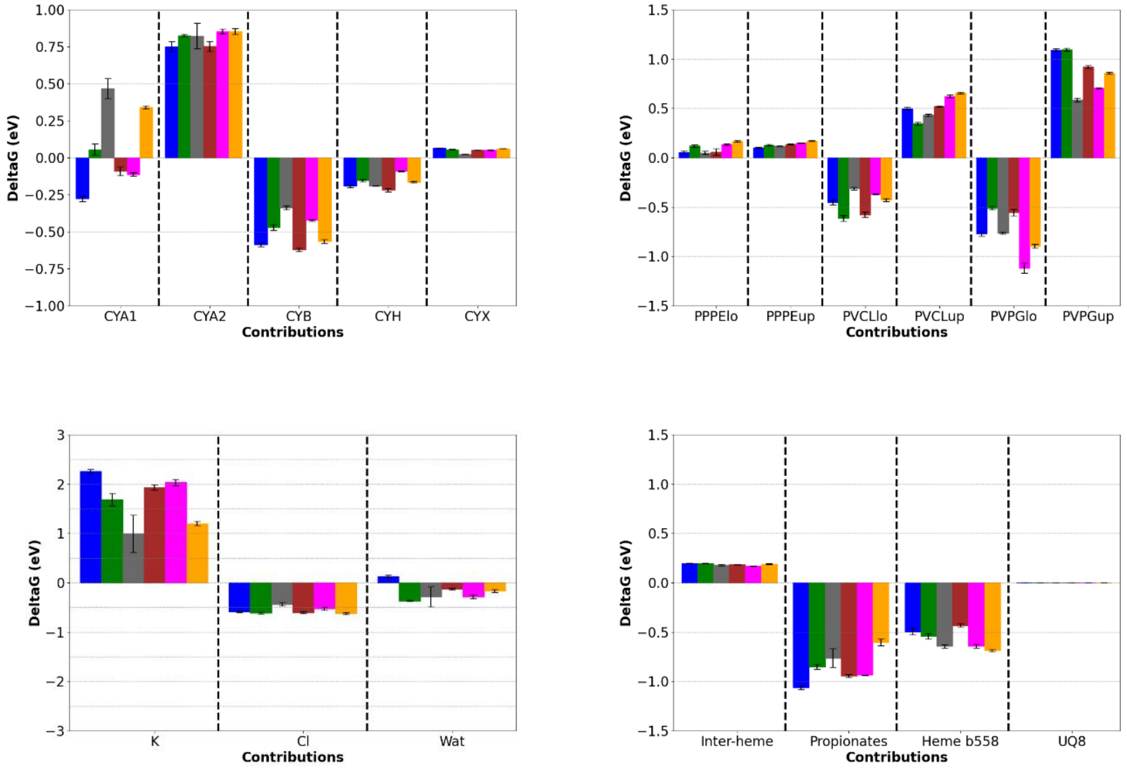


*Figure S8: Decomposition of the outer-sphere electron transfer free energy of $b_{595}$ to heme d electron transfer into various contributions. Top-left: contributions of each proteic chain; top-right: decomposition of the contribution of the lipids with respect to the type of lipid (PPPE, PVCL2 and PVPG) and the leaflet (lo for lower and up for upper); bottom-left: water and counter-ion contributions; bottom-right: contribution of the heme $b_{595}$/heme d interaction (without taking into account the propionates), of the propionates of hemes $b_{595}$ and heme d, of heme $b_{558}$, and of the UQ8 quinone. Setup SA1: blue, setup SA2: green, setup SB1: grey, setup SB2: brown, setup SC1: magenta, setup SC2: orange.*

*Table S1: DFT calculations for heme cofactors in the gas phase. ΔE are relative energies between redox and spin states for a given complex. $IP_v$ and $EA_v$ are the vertical ionization potential and electron affinity, the spin state of the resulting species being given in bracket (in eV); s, t qi, d, q and se respectively stand for singlet, triplet, quintet, doublet, quartet and sextet states. d(Fe-Xap) denotes the iron-ligand distances in Å, and $\lambda_{1/2}$ are the internal reorganization energies (see main text). All values are those obtained with the ωB97X-D3 functionals on the geometries optimized with PBE functional.*

| | $\Delta E$ | $IP_v/EA_v$ | d(Fe-Xap) | $\lambda_{1/2}$ |
|---|---|---|---|---|
| heme *b*$_{558}$ | | | | |
| $Fe^{2+}$ *singlet* | 0.00 | 5.38 (d) | 2.88 ($S_{met}$)/2.21 ($N_{his}$) | 0.10 |
| $Fe^{2+}$ *triplet* | 0.30 | 5.67 (d) | 2.30 ($S_{met}$)/1.96 ($N_{his}$) | 0.69 |
| | | 5.18 (q) | | 0.15 |
| $Fe^{3+}$ *doublet* | 5.28 | 5.26 (s) | 2.33 ($S_{met}$)/1.97 ($N_{his}$) | 0.02 |
| | | 4.25 (t) | | 0.73 |
| $Fe^{3+}$ *quartet* | 5.33 | 5.03 (t) | 2.78 ($S_{met}$)/2.21 ($N_{his}$) | 0.01 |
| heme *b*$_{595}$ | | | | |

| | | | | |
|---|---|---|---|---|
| $Fe^{2+}$ *singlet* | 0.91 | 5.77 (d) | 1.88 ($O_{glu}$) | 0.33 |
| $Fe^{2+}$ *triplet* | 0.00 | 6.30 (d) | 2.29 ($O_{glu}$) | -0.04 |
| | | 5.67 (q) | | 0.10 |
| $Fe^{2+}$ *quintet* | 0.10 | 5.76 (q) | 2.16 ($O_{glu}$) | 0.29 |
| | | 6.12 (se) | | 0.28 |
| $Fe^{3+}$ *doublet* | 6.34 | 5.51 (s) | 1.95 ($O_{glu}$) | -0.07 |
| | | 5.87 (t) | | 0.47 |
| $Fe^{3+}$ *quartet* | 5.57 | 5.40 (t) | 2.17 ($O_{glu}$) | 0.17 |
| $Fe^{3+}$ *sextet* | 5.94 | 5.71 (qi) | 2.11 ($O_{glu}$) | 0.13 |
| $H_2O$-Fe2$^+$ *singlet* | 0.00 | 5.56 (d) | 1.88 ($O_{glu}$)/2.04 ($O_{H2O}$) | 0.06 |
| $H_2O$-$Fe^{2+}$ *triplet* | 0.00 | 6.38 (d) | 2.30 ($O_{glu}$)/3.31 ($O_{H2O}$) | 0.88 |
| | | 5.79 (q) | | 0.68 |
| $H_2O$-$Fe^{2+}$ *quintet* | 0.05 | 5.92 (q) | 2.19 ($O_{glu}$)/3.32 ($O_{H2O}$) | 0.86 |
| | | 6.26 (se) | | 0.91 |
| $H_2O$-$Fe^{3+}$ *doublet* | 5.50 | 5.37 (s) | 1.94 ($O_{glu}$)/1.99 ($O_{H2O}$) | 0.13 |
| | | 4.16 (t) | | 1.34 |
| $H_2O$-$Fe^{3+}$ *quartet* | 5.12 | 5.22 (t) | 2.29 ($O_{glu}$)/2.44 ($O_{H2O}$) | -0.10 |
| $H_2O$-$Fe^{3+}$ *sextet* | 5.40 | 5.52 (qi) | 2.21 ($O_{glu}$)/2.36 ($O_{H2O}$) | -0.17 |
| heme *d* | | | | |
| $Fe^{2+}$ *singlet* | 0.26 | 5.74 (d) | 1.86 ($N_{his}$) | 0.06 |
| $Fe^{2+}$ *triplet* | 0.00 | 6.12 (d) | 2.11 ($N_{his}$) | 0.18 |
| | | 5.54 (q) | | 0.24 |
| $Fe^{2+}$ *quintet* | 0.43 | 4.85 (q) | 2.10 ($N_{his}$) | 0.24 |
| | | 5.95 (se) | | 1.08 |
| $Fe^{3+}$ *doublet* | 5.68 | 5.56 (s) | 1.88 ($N_{his}$) | 0.12 |
| | | 5.34 (t) | | 0.61 |
| $Fe^{3+}$ *quartet* | 5.03 | 5.24 (t) | 2.09 ($N_{his}$) | 0.06 |
| $Fe^{3+}$ *sextet* | 5.30 | 5.55 (qi) | 2.05 ($N_{his}$) | -0.68 |
| $H_2O$-Fe2$^+$ *singlet* | 0.00 | 5.47 (d) | 1.88 ($N_{his}$)/2.04 ($O_{H2O}$) | 0.03 |
| $H_2O$-$Fe^{2+}$ *triplet* | 0.42 | 6.01 (d) | 2.30 ($N_{his}$)/3.31 ($O_{H2O}$) | 0.99 |
| | | 5.69 (q) | | 0.77 |
| $H_2O$-$Fe^{2+}$ *quintet* | 1.10 | 4.97 (q) | 2.11 ($N_{his}$)/3.43 ($O_{H2O}$) | 0.72 |
| | | 5.53 (se) | | -0.08 |
| $H_2O$-$Fe^{3+}$ *doublet* | 5.44 | 5.31 (s) | 2.04 ($N_{his}$)/1.92 ($O_{H2O}$) | 0.13 |
| | | 4.40 (t) | | 0.62 |
| $H_2O$-$Fe^{3+}$ *quartet* | 5.35 | 4.99 (t) | 2.13 ($N_{his}$)/2.49 ($O_{H2O}$) | -0.06 |
| $H_2O$-$Fe^{3+}$ *sextet* | 6.71 | 5.49 (qi) | 2.10 ($N_{his}$)/3.48 ($O_{H2O}$) | 0.11 |

*Table S2: Decomposition of the free energy of $b_{595}$ to heme d electron transfer corresponding to figure 2 of the main text.*

| Outer sphere contribution | $\Delta G$ (eV) | $\Delta G_{prot}$ (eV) | $\Delta G_{lipid}$ (eV) | $\Delta G_{wations}$ (eV) | $\Delta G_{other}$ (eV) |
|---|---|---|---|---|---|
| SA1 | 0.72 (0.02) | -0.24 (0.03) | 0.52 (0.04) | 1.81 (0.04) | -1.56 (0.02) |
| SA2 | 0.38 (0.05) | 0.31 (0.04) | 0.57 (0.03) | 0.7 (0.14) | -1.4 (0.05) |
| SB1 | -0.05 (0.07) | 0.79 (0.02) | 0.11 (0.04) | 0.28 (0.16) | -1.41 (0.11) |
| SB2 | 0.39 (0.02) | -0.13 (0.08) | 0.51 (0.05) | 1.2 (0.05) | -1.38 (0.03) |
| SC1 | 0.22 (0.03) | 0.28 (0.02) | 0.12 (0.04) | 1.23 (0.09) | -1.58 (0.02) |
| SC2 | 0.36 (0.02) | 0.53 (0.03) | 0.52 (0.04) | 0.41 (0.06) | -1.29 (0.04) |

*Table S3: Decomposition of the reorganization free energy of $b_{595}$ to heme d electron transfer. Other include heme b558, propionate groups, UQ8 and inter-heme interaction*

| Outer sphere contribution | $\lambda$ (eV) | $\lambda_{prot}$ (eV) | $\lambda_{lipid}$ (eV) | $\lambda_{wations}$ (eV) | $\lambda_{other}$ (eV) |
|---|---|---|---|---|---|
| SA1 | 1.22 (0.01) | 0.45 (0.02) | -0.03 (0.02) | 0.78 (0.03) | 0.04 (0.01) |
| SA2 | 1.05 (0.03) | 0.77 (0.02) | 0.01 (0.04) | 0.10 (0.04) | 0.17 (0.03) |
| SB1 | 1.02 (0.03) | 1.15 (0.02) | 0.10 (0.08) | -0.11 (0.12) | -0.12 (0.04) |
| SB2 | 1.00 (0.01) | 0.57 (0.02) | 0.11 (0.05) | 0.09 (0.06) | 0.25 (0.01) |
| SC1 | 1.03 (0.02) | 0.76 (0.01) | -0.09 (0.05) | 0.44 (0.03) | -0.09 (0.01) |
| SC2 | 1.33 (0.01) | 0.64 (0.03) | 0.17 (0.02) | 0.73 (0.03) | -0.20 (0.03) |

*Table S4: Decomposition of the free energy of $b_{595}$ to heme d electron transfer for protein components.*

| Outer sphere contribution | $\Delta G_{CYA1}$ (eV) | $\Delta G_{CYA2}$ (eV) | $\Delta G_{CYB}$ (eV) | $\Delta G_{CYH}$ (eV) | $\Delta G_{CYX}$ (eV) |
|---|---|---|---|---|---|
| SA1 | -0.28 (0.02) | 0.75 (0.03) | -0.59 (0.01) | -0.19 (0.01) | 0.06 (0.0) |
| SA2 | 0.06 (0.04) | 0.83 (0.01) | -0.47 (0.02) | -0.15 (0.01) | 0.06 (0.0) |
| SB1 | 0.47 (0.07) | 0.82 (0.09) | -0.34 (0.01) | -0.19 (0.0) | 0.02 (0.0) |
| SB2 | -0.09 (0.03) | 0.75 (0.03) | -0.62 (0.01) | -0.22 (0.01) | 0.05 (0.0) |
| SC1 | -0.11 (0.01) | 0.85 (0.02) | -0.42 (0.01) | -0.09 (0.0) | 0.05 (0.0) |
| SC2 | 0.34 (0.01) | 0.85 (0.02) | -0.56 (0.01) | -0.16 (0.01) | 0.06 (0.0) |

*Table S5: Decomposition of the reorganization free energy of $b_{595}$ to heme d electron transfer for protein components.*

| Outer sphere contribution | $\lambda_{CYA1}$ (eV) | $\lambda_{CYA2}$ (eV) | $\lambda_{CYB}$ (eV) | $\lambda_{CYH}$ (eV) | $\lambda_{CYX}$ (eV) |
|---|---|---|---|---|---|
| SA1 | 0.38 (0.01) | -0.01 (0.01) | 0.03 (0.01) | 0.04 (0.01) | 0.00 (0.01) |
| SA2 | 0.64 (0.02) | 0.06 (0.02) | 0.09 (0.01) | -0.01 (0.01) | -0.01 (0.01) |
| SB1 | 0.96 (0.03) | 0.22 (0.05) | 0.01 (0.01) | -0.02 (0.01) | -0.01 (0.01) |
| SB2 | 0.59 (0.01) | -0.01 (0.01) | 0.01 (0.01) | -0.02 (0.01) | -0.01 (0.01) |
| SC1 | 0.60 (0.01) | 0.10 (0.01) | 0.05 (0.01) | 0.01 (0.01) | 0.00 (0.01) |
| SC2 | 0.60 (0.02) | 0.06 (0.02) | -0.04 (0.01) | 0.00 (0.01) | 0.01 (0.01) |

*Table S6: Decomposition of the free energy of $b_{595}$ to heme d electron transfer for lipid components.*

| Outer sphere contribution | $\Delta G_{PPPElo}$ (eV) | $\Delta G_{PPPEup}$ (eV) | $\Delta G_{PVCLlo}$ (eV) | $\Delta G_{PVCLup}$ (eV) | $\Delta G_{PVPGlo}$ (eV) | $\Delta G_{PVPGup}$ (eV) |
|---|---|---|---|---|---|---|
| SA1 | 0.06 (0.01) | 0.1 (0.01) | -0.45 (0.03) | 0.5 (0.01) | -0.77 (0.02) | 1.09 (0.01) |
| SA2 | 0.12 (0.01) | 0.13 (0.01) | -0.61 (0.03) | 0.35 (0.01) | -0.51 (0.02) | 1.1 (0.01) |
| SB1 | 0.05 (0.02) | 0.12 (0.0) | -0.31 (0.01) | 0.43 (0.01) | -0.77 (0.01) | 0.59 (0.02) |
| SB2 | 0.06 (0.03) | 0.14 (0.01) | -0.58 (0.03) | 0.52 (0.01) | -0.55 (0.03) | 0.92 (0.01) |
| SC1 | 0.13 (0.01) | 0.15 (0.0) | -0.37 (0.01) | 0.62 (0.01) | -1.12 (0.05) | 0.7 (0.01) |
| SC2 | 0.17 (0.01) | 0.17 (0.0) | -0.43 (0.01) | 0.66 (0.01) | -0.9 (0.02) | 0.86 (0.01) |

*Table S7: Decomposition of the reorganization free energy of $b_{595}$ to heme d electron transfer for lipid components.*

| Outer sphere contribution | $\lambda_{PPPElo}$ (eV) | $\lambda_{PPPEup}$ (eV) | $\lambda_{PVCLlo}$ (eV) | $\lambda_{PVCLup}$ (eV) | $\lambda_{PVPGlo}$ (eV) | $\lambda_{PVPGup}$ (eV) |
|---|---|---|---|---|---|---|
| SA1 | 0.04 (0.01) | 0.01 (0.01) | 0.05 (0.01) | -0.05 (0.01) | -0.06 (0.02) | -0.02 (0.01) |
| SA2 | -0.04 (0.01) | 0.01 (0.01) | 0.01 (0.01) | -0.02 (0.01) | 0.13 (0.01) | -0.07 (0.01) |
| SB1 | 0.01 (0.01) | -0.02 (0.01) | 0.15 (0.01) | -0.06 (0.01) | 0.08 (0.04) | -0.06 (0.02) |
| SB2 | 0.02 (0.03) | -0.01 (0.01) | -0.04 (0.02) | 0.04 (0.01) | 0.14 (0.03) | -0.04 (0.01) |
| SC1 | 0.01 (0.01) | 0.03 (0.01) | 0.02 (0.01) | -0.05 (0.02) | 0.02 (0.02) | -0.12 (0.01) |
| SC2 | -0.01 (0.01) | 0.01 (0.01) | 0.05 (0.02) | 0.04 (0.01) | -0.01 (0.03) | 0.09 (0.02) |

*Table S8: Decomposition of the free energy of $b_{595}$ to heme d electron transfer for water and counterions.*

| Outer sphere contribution | $\Delta G_K$ (eV) | $\Delta G_{Cl}$ (eV) | $\Delta G_{wat}$ (eV) |
|---|---|---|---|
| SA1 | 2.26 (0.05) | -0.58 (0.02) | 0.13 (0.03) |
| SA2 | 1.69 (0.13) | -0.62 (0.02) | -0.37 (0.01) |
| SB1 | 1.0 (0.38) | -0.43 (0.04) | -0.28 (0.2) |
| SB2 | 1.93 (0.05) | -0.6 (0.02) | -0.13 (0.02) |
| SC1 | 2.03 (0.06) | -0.52 (0.03) | -0.28 (0.04) |
| SC2 | 1.2 (0.05) | -0.63 (0.02) | -0.16 (0.03) |

*Table S9: Decomposition of the reorganization free energy of $b_{595}$ to heme d electron transfer for water and counterions.*

| Outer sphere contribution | $\lambda_K$ (eV) | $\lambda_{Cl}$ (eV) | $\lambda_{wat}$ (eV) |
|---|---|---|---|
| SA1 | 0.32 (0.03) | 0.07 (0.02) | 0.38 (0.04) |
| SA2 | -0.07 (0.02) | -0.07 (0.01) | 0.24 (0.04) |
| SB1 | 0.13 (0.23) | 0.05 (0.03) | -0.30 (0.14) |
| SB2 | -0.03 (0.04) | -0.04 (0.01) | 0.16 (0.02) |
| SC1 | 0.05 (0.04) | 0.11 (0.01) | 0.29 (0.03) |
| SC2 | 0.70 (0.07) | -0.09 (0.01) | 0.13 (0.04) |

*Table S10: Decomposition of the free energy of $b_{595}$ to heme d electron transfer for other components*

| Outer sphere contribution | $\Delta G_{inter-heme}$ (eV) | $\Delta G_{prop}$ (eV) | $\Delta G_{b558}$ (eV) | $\Delta G_{UQ8}$ (eV) |
|---|---|---|---|---|
| SA1 | 0.2 (0.0) | -1.06 (0.02) | -0.49 (0.03) | -0.0 (0.0) |
| SA2 | 0.2 (0.0) | -0.85 (0.02) | -0.54 (0.03) | 0.0 (0.0) |
| SB1 | 0.18 (0.01) | -0.76 (0.1) | -0.65 (0.02) | 0.0 (0.0) |
| SB2 | 0.18 (0.0) | -0.94 (0.01) | -0.43 (0.02) | 0.0 (0.0) |
| SC1 | 0.17 (0.0) | -0.94 (0.0) | -0.64 (0.02) | 0.0 (0.0) |
| SC2 | 0.19 (0.0) | -0.61 (0.03) | -0.69 (0.01) | 0.0 (0.0) |

*Table S11: Decomposition of the reorganization free energy of $b_{595}$ to heme d electron transfer for other components*

| Outer sphere contribution | $\lambda_{inter-heme}$ (eV) | $\lambda_{prop}$ (eV) | $\lambda_{b558}$ (eV) | $\lambda_{UQ8}$ (eV) |
|---|---|---|---|---|
| SA1 | -0.02 (0.01) | 0.06 (0.01) | -0.02 (0.01) | 0.00 (0.00) |
| SA2 | 0.00 (0.01) | 0.10 (0.01) | 0.08 (0.02) | 0.00 (0.00) |
| SB1 | -0.01 (0.01) | -0.06 (0.03) | -0.06 (0.01) | 0.00 (0.00) |
| SB2 | -0.02 (0.01) | 0.19 (0.02) | 0.06 (0.01) | 0.00 (0.00) |
| SC1 | 0.01 (0.01) | -0.03 (0.01) | -0.06 (0.01) | 0.00 (0.00) |
| SC2 | -0.01 (0.01) | -0.11 (0.03) | -0.09 (0.01) | 0.00 (0.00) |
